## Appendix S1 for "Predicting direct and indirect non-target impacts of biocontrol agents using machine-learning approaches"

#### Traits and phylogenies

##### Body size

Body size is correlated with a number of important characteristics [1], and is often the best predictor of species interactions (see references in main text). We therefore measured body mass of both host and parasitoid species. Lepidoptera larvae ('hosts') were weighed immediately after collection (i.e. fresh weight), and dry weight was measured for parasitoids. Because parasitoids were stored in ethanol 70% after their emergence, to remove all the water content in their bodies we sequentially moved them to different Eppendorf tubes of increasing ethanol concentrations (80%, 90%, 95% and twice in 100%). We then removed the parasitoids from the 100% ethanol concentration Eppendorf tubes and put them in a vacuum desiccator to extract all the liquid from them. We left the parasitoids in the desiccator for 24 hours and immediately after removing them from the desiccator we weighed them to obtain their dry weight. Body mass was averaged at species level, except for 3 parasitoid and 13 host species, which were measured at genus (or family) level (i.e. the average of all species in the same genus (or family) was used). Species level estimates were not available for these species because the 13 host species could only be identified when they developed into adults and the 3 parasitoid species had already been deposited at the New Zealand Arthropod Collection (NZAC) in Auckland and the Te Papa Museum Entomology Collection in Wellington, New Zealand. Additionally, body mass of 2 host and 2 parasitoid species were not available even at family level (for the same reasons described above), so the average body mass of all hosts and parasitoids, respectively, were used for these species.

##### Generality

We calculated host and parasitoid generality, because recent evidence suggests that species generalism tends to be conserved when a species moves to a novel range [2], such as in species invasion or biocontrol introductions, so it may be a useful characteristic to include when predicting interactions.

As a measure of species trophic generalism, we used normalised degree (ND), which represents the number of species with which a given species interacts, normalised to account for differences in the availability of interaction partners [3], and thus is a measure of the relative generality of a species compared to other species in the same trophic level, within the same network [3]. We calculated the ND of host and parasitoid species using the "specieslevel" function of the bipartite v. 2.15 R package [4], to represent how specialist/generalist host and parasitoid species were in their interactions with plant and host species, respectively.

ND was calculated separately for each forest type (native vs. plantation) because trophic generality can change across habitats [5], and was calculated using only the training data (i.e. the generality of species at the test sites was estimated by their generality at the training sites). This is important because parasitoid ND is calculated from the host-parasitoid network and it would have been circular to use information from the test network to predict the test network. Species that were only present at the test sites would be the equivalent of novel species such as introduced biocontrol agents for which there are no existing data on generality. For these species, genus level (or family level) averages of ND were used instead of species level estimates (i.e. the average ND value of all species in the same genus (or family) in the training data was used). This could be a reasonable starting approximation, because species' network roles can be conserved at the family or higher level [6]. For two parasitoid species, no species in the same family were present in the training data, so the average ND value of all parasitoid species was used.

The generality of a species often appears to increase with its abundance [7, 8], in part because a larger sample size characterises a greater proportion of possible interactions. Thus, when predicting interactions, we included abundance data (see below) separately from generality, and therefore did not control for abundance when calculating species generality.

#### Biogeographic status

We included biogeographic status (native vs. exotic) of host species as a trait because recent evidence suggests that niche processes for species interactions (and therefore predictions based on traits and phylogenies) are weaker for exotic species [9]. Host biogeographic status was determined using Hoare *et al.* [10], Hoare [11] and Philpott [12]. We were not able to determine the biogeographic status of 7 host species (Psychidae 4 spots, Psychidae other, Psychidae sp2, Strepsicrates sp, Psychidae, Stathmopoda sp chocolate, and Archipini sp) and therefore removed these species from all subsequent analyses. We did not include biogeographic status for parasitoid species because we were not able to determine this information for parasitoid species identified to morphospecies or genus level (since to our knowledge none of the genera present in our dataset are known to be entirely native (or exotic)).

#### Abundance and phenology

Species abundance and phenological overlap play a key role in structuring interactions [13-18]. We therefore measured phenology and abundance (for both host and parasitoid species). We measured phenology and abundance separately for each forest type, and separately for training and testing sites. We considered two aspects of phenology: abundance of each species in each month and phenological overlap between each host-parasitoid pair.

##### Abundance

Site level abundance at the test sites was calculated as the abundance of each species at each site summed across both sampling months in the 'before' time step (*t*, October, November 2010) and divided by two (to obtain the abundance of each species per site per month, so that the values are comparable to the training data). To calculate site level abundance for the training data, we used the abundance of each species summed across all training sites and sampling months averaged by the number of sites and months to account for sample size differences across the training and testing datasets. We summed abundance in this way because the training data comprise one meta-web for each random-forest and KNN model; see 'Training meta-networks' below.

#### Phenology

We measured phenology as the abundance of each species in each month. To calculate this aspect of phenology, we first estimated the abundance of each host species as the number of individuals collected each month. No independent estimates of adult parasitoid abundance were available, so we estimated the abundance of each parasitoid species from the quantitative interaction networks by summing across link weights (representing parasitism rates, i.e. the abundance of parasitoid larvae in hosts) across sites. Importantly, although parasitoid abundance was calculated from the network, interaction data (i.e., links between species) were not used for calculating abundance, avoiding the circularity of using the network to predict the network. However, we acknowledge that this requires knowledge of the abundance of each parasitoid within the community for which the predictions are being made. Nevertheless, if successful as a proof of concept, this approach has the advantage of being able to test (using simulation) how impacts may change with increasing abundance of a parasitoid species, though we do not focus on this here as we lack the data to validate such simulations. We normalised abundance of host and parasitoid species by the number of sites sampled per month. We used two sampling months (October and November 2010) for phenology, as these were the only sampling months in common between the training and testing datasets.

We calculated the quantitative phenological overlap for each host-parasitoid species pair following the method described in Pleasants [19]. We first divided the abundance of each species in each month by the total abundance of that species across all months, to obtain the proportional abundance of each species per month. Then, for each host-parasitoid pair, we compared, month-by-month, the proportional abundance of both the host and parasitoid species and took the lower value of the two (because the abundance of the rarer partner limits the interaction frequency). This gives the 'overlap' for each pair for each month. We then summed these overlap values to obtain the final phenological overlap value for each host-parasitoid pair. For the training data, all 7 sampling months were used to calculate phenological overlap, while for the test data, two sampling months (October and November 2010) were used to calculate phenological overlap.

### Phylogenies

Phylogenies, the shared evolutionary history of species, can serve as a surrogate for variation in many traits (both known and unknown), and can therefore help explain network structure [20]. Additionally, due to convergent evolution, traits are not always perfectly correlated with phylogenies, so phylogenies can provide additional information about coevolutionary history. Unfortunately, highly-resolved phylogenies are not always available at the species level, which could hinder their widespread use in biocontrol risk assessment. One potential solution would be taxonomic trees, though these lose information about separation times of taxa and thereby generate 'soft polytomies' (e.g., all species within a genus appear to have branched at the same time and thus have equal phylogenetic distance). As a compromise between the need for more detail than taxonomic trees, but without depending on well-resolved phylogenies, we constructed both host and parasitoid phylogenies using publicly-available data on two marker sequences (see below). Our aim here was to have an easily-repeatable method to capture phylogenetic relationships among species, rather than to create a well-resolved phylogeny as a goal in itself. Accordingly, we chose this approach of constructing phylogenies using a small number of sequences (two) as they provide more information than taxonomic trees, while not requiring so many sequences that availability would become a constraint.

#### Host phylogeny

We constructed a host phylogeny for the host species in the dataset (see 'Study system' in Methods in Main text) using one mitochondrial marker (Cytochrome Oxidase subunit 1, COI) and one nuclear marker (wingless gene, Wgl) sequence, both of which we obtained from GenBank [21]. We chose these sequences because they were available for the greatest number of species in our dataset. We used the rentrez v 1.2.2 R package [22] to obtain GenBank accession numbers for both markers for each host species. We then used the ape v 5.4 R package [23] to retrieve sequences from GenBank using this list of accession numbers.

The Wgl marker sequence was used at the family level (i.e. the same Wgl sequence was used for all genera within the same family) to create a backbone phylogeny at the family level, and the COI marker was used at species level where available, and at genus or family level where a species level sequence were not available (i.e. for species that did not have a GenBank entry for a sequence, the sequence from another species within the same genus or family was used instead). Note that this genus- or family- level sequence was not the same as any species-level sequence we may have had for other species in our dataset, except for the genus Tatosoma where only one sequence was available in GenBank and was therefore assigned to all species within the genus. When this case of species- (or genus-) level sequences not being available applied to more than one species within the same genus (or more than one genus within the same family) in our dataset, we included only one species from such a set of species (within the same genus or family) in the phylogeny and assigned the same axis values to all species in the set (see 'Including phenology and phylogenies in models' below). In other words, congeneric species (or species in the same family) could not be distinguished phylogenetically when neither one occurred in GenBank for the COI sequence. Of the 45 genera present in our dataset, 9 genera contained at least two species (in our dataset) that did not occur in GenBank for a species-level COI sequence, and for an additional 10 genera, only family-level COI sequences were available. We did this assignment of the same axis values to all species in a set rather than assigning the same GenBank accession number, and therefore sequence, to each of them as this would have resulted in including multiple species in the phylogeny with identical sequences for both markers.

The host phylogeny was constructed using 40 host species in the dataset. Although our host phylogeny does not include all host species in our dataset (discussed above), all families in our dataset are represented in the phylogeny. The number of COI sequences available at family, genus and species level were 4, 20, 15 respectively (for one species we did not include a COI sequence; see below).

The COI and WgI sequences were aligned separately in MEGA X (Molecular Evolutionary Genetics Analysis) [24] (v 10.1.8), using MUSCLE [25] with default settings. The COI sequences for two species (*Heterocrossa sp indet C* and *Graphania chlorodonta*) were removed from the alignment as they did not align well with the other sequences. Because *Heterocrossa sp indet C* was the only species (in our dataset) in the Carposinidae family, we included this species in the phylogeny using a dummy sequence of N's for the COI sequence, as the WgI family sequence would be able to distinguish this species (and its unique family) from the other species. However, we removed *Graphania chlorodonta* from the phylogeny as, without a COI sequence, it would not be able to be distinguished from the other species in the same family (Noctuidae). We assigned the average axis value of all species within the family Noctuidae (that were present in our phylogeny) to *Graphania chlorodonta* (see 'Including phenology and phylogenies in models' below). Note that a genus level average was not available for this species because a COI sequence in GenBank did not occur for any species within the genus *Graphania*. To incorporate both markers, we imported both aligned sequence files into BEAUTi2 (v 2.6.2) (available within the BEAST2 package) which is equivalent to concatenating the sequences (i.e. each host species has a concatenated two-gene sequence (COI + WgI)). We included *Pogonomyrmex subdentatus* (Hymenoptera: Formicidae) as an outgroup. The outgroup was included as a prior with a lognormal distribution (mean = 10, standard deviation = 0.5, offset = 340).

BEAST2 (v 2.6.2) was used to estimate the phylogenies [26, 27]. The tree prior was set using a Yule model (this is the same model used by Peralta *et al.* [28] to construct phylogenies for the same taxa).

We linked the tree in BEAUTi2 to ensure the same phylogeny was used for all partitions (sequences) and linked the clock model (which assumes that the partitions have the same evolutionary branch-rate distribution). We used a GTR site model [29] with a Gamma Category Count of 4 for both partitions (again this is the same model used by Peralta *et al.* [28] to construct phylogenies for the same taxa). We used a lognormal relaxed molecular clock, and the chain length was set to 10,000,000, sampling parameters every 1000 trees. We used Tree Annotator (v 2.6.2) (available within the BEAST2 package) to construct a maximum credibility tree, with a burn-in of 1000 initial trees, and FigTree (v 1.4.4) [30] was used to view the phylogenetic tree (Fig 1A in Appendix S1).

**Fig 1 in Appendix S1. (A) Host and (B) parasitoid phylogenies, with phylogenetic relationships inferred using DNA sequences obtained from GenBank.**

(A) Host phylogeny with phylogenetic relationships based on one nuclear marker (Wgl) and one mitochondrial marker (COI). We included *Pogonomyrmex subdentatus* (Hymenoptera: Formicidae) as an outgroup, because all of the host species in our dataset were from the same order (Lepidoptera). (B) Parasitoid phylogeny with phylogenetic relationships based on one ribosomal marker (28S) and one mitochondrial marker (COI). We included *Sitona discoideus* (Coleoptera: Curculionidae) as the outgroup, because the parasitoids in our data set belonged to the orders of Hymenoptera and Diptera. Because we used COI sequences at genus or family level where a species-level sequence was not available, some genera are split across branches. However, all genera within the same family occur on the same branch.

#### Parasitoid phylogeny

We constructed a parasitoid phylogeny (for  the parasitoid species in the dataset; see 'Study system' in Methods in Main text) using one mitochondrial marker (COI) and one ribosomal marker (28S), both obtained from GenBank [21]. Frost *et al.* [31] sequenced COI for some parasitoid species in our dataset (GenBank accession numbers: KM106857 – KM107193). We used these sequences and searched GenBank for species that Frost *et al.* [31] did not sequence. We constructed the phylogeny using the

28S marker sequence at the family level, and the COI sequence at the species level (or at genus or family level when a species level sequence was not available). As with the host species (see above), COI sequences were not available for every parasitoid species; we therefore constructed the parasitoid phylogeny using 49 of the 68 parasitoid species in our dataset. The number of COI sequences available at family, genus and species (or morpho-species; i.e. sequenced by [31]) level were 3, 13 and 33, respectively. We used the same phylogenetic construction method as explained for hosts, but this time *Sitona discoideus* (Coleoptera: Curculionidae) was included as the outgroup (Fig 1B in Appendix S1).

### Machine-learning techniques

#### Random forest

Random forest is a machine learning technique which can be used for both regression and classification problems. A random forest is constructed from many (often hundreds or thousands) individual decision trees. Individual estimates from each tree are combined to give an overall prediction (average or majority vote for regression and classification, respectively). Each tree is trained on a random sample of the data, and only a random subset of features (i.e. predictor variables) are considered for the splitting of each node. This is important because it ensures that trees are independent (not correlated) and diverse (i.e. use different variables to make predictions).

#### KNN (K Nearest Neighbours)

Recommender systems are a class of methods used to predict the rating or preference that a user would give to a presently unknown item, based on their ratings of known items (or based on ratings of similar users). They are widely used by companies such as YouTube, Netflix and Spotify for generating playlists, and by Amazon (and other online services) for recommending items to users. KNN is one such recommender system, which uses the preferences of similar users to recommend new items (such as films) to users. KNN is based on the idea that similar users will prefer similar items. More recently, KNN has been used to predict interactions in ecological communities [32, 33].

We used traits and phylogenies to determine similarity between parasitoid species. As a measure of which species had (dis)similar traits and phylogenetic position, we calculated the Euclidean distance between parasitoid species in trait-and-phylogenetic space using the 'distance matrix' function from the SciPy Spatial v 1.4.1 package [34]. We used Euclidean distance because parasitoid phylogeny (included as PCOA axes; see below) and all parasitoid traits were continuous. We scaled parasitoid phylogeny and all traits to have a range of 0 to 1 (using the MinMaxScaler function from the scikit-learn v 0.23.1 package [35]) and weighted the individual columns of phenology and phylogeny by half (as these characteristics each consisted of two components; first and second PCOA axes, and abundance in October and November 2010, respectively) so that these characteristics would each have a weighting equal to other traits. We scaled all traits for the training and testing data together, prior to any hyper-parameter tuning or model fitting processes, so that the scaled trait values for species within the training and testing data were comparable (i.e. on the same scale).

#### Including phenology and phylogenies in models

We included two aspects of phenology (abundance of each species in each month and phenological overlap between each host-parasitoid pair; see 'Abundance and phenology' above) in the random-forest models. We did not include host-parasitoid phenological overlap in the KNN models, as KNN considers host and parasitoid traits separately, in contrast to random forest which considers host-parasitoid trait matching.

Phylogenetic relationships between species were included in models by first converting the phylogenetic tree into a distance matrix (using the cophenetic function in the ape v 5.4 R package [23]) and then converting the distance matrix into principal coordinate axes (using the pcoa function in the ape package), to include in the machine-learning models. The first and second PCOA axes explained 33.78 and 21.38 percent of the variation, respectively, for the host phylogeny and 42.96 and 25.88 percent of the variation, respectively, for the parasitoid phylogeny. The third PCOA axis explained less variance for both phylogenies (12.16 and 8.72 percent, respectively). We therefore only included the first two PCOA axes.

#### Comparison of data requirements for random forest and KNN

Random forest and KNN both require known interactions to train the model to predict interactions at a new location or for new species (e.g., control agents). In addition to known interactions, trait data can be used by both random forest and KNN to predict interactions. Our data was structured so that a row of data included the host and parasitoid identity, whether or not they interacted, along with parasitoid traits and host traits for random forest, and just parasitoid traits for KNN. In other words, each row of our data-frame was a host-parasitoid species pair. However, for random forest, a model can be trained without knowing the identities of the species present in the training network (i.e. the known interactions used to create the model), as only their traits (and not species' identities) are used as predictor variables. This would be an advantage in cases where interaction and trait data have been collected, but species have not been identified (or have only been identified to genus or family level). In this case, a random-forest model could be trained on individual level network and trait data, where links are recorded between individuals rather than species (e.g., [36]). Individual network models have the advantage of being able to predict interactions among species that, for example, have large ontogenetic changes in diet (e.g., [37]).

KNN (and other recommender systems that use the preferences of similar users to recommend new items to users) must determine the similarity between parasitoids (users) (and hosts (items) in the case of 'new hosts'; see 'KNN – adding 'new' host species' below) in order to make predictions. Traits can be used to determine similarity between species, and in the case of 'new host species' (which occur in the testing but not training data), both host and parasitoid traits would be required (akin to random forest). However, in order to determine the likely hosts of a given parasitoid species, where all potential host species were present in the training data, only parasitoid (and not host) trait data would be required: host traits are only required to predict interaction partners for new hosts. Furthermore, if trait data were not available, interaction data could be used to infer similarity between species, though only if they overlap in some hosts (for example, parasitoid species that interact with similar host species are considered similar). Combined, these characteristics of recommender systems mean that KNN could potentially be used to predict potential host species of new parasitoid species (e.g. control agents) using only known interaction preferences of similar parasitoid species (i.e. parasitoid species with similar host-use; see above), without any trait data being required, such as in regions with poorly-studied fauna, allowing all data to be used. Therefore, in terms of data requirements, KNN is more flexible than random forest. We only focussed on using traits (rather than interactions) to determine the similarity between parasitoid species, as it is more likely to have trait data available for a proposed agent.

### Training and testing machine-learning models

#### Training meta-networks

The training meta-networks were the data on host-parasitoid combinations (for which an interaction occurred or not), pooled across all training sites, that were used to train the KNN and random forest models. As described in the main text, KNN models were trained on weighted meta-networks and random-forest models were trained on binary meta-networks. They only included host species that were parasitized at least once in the dataset. Further, we only included a host-parasitoid species pair as non-interacting if they co-occurred at least once at the same site, as host-parasitoid pairs that did not co-occur cannot be assumed to not interact in situations where they are both present. In addition to this, networks of species interactions are often sparse (there are few interactions compared to non-interactions). This can cause random forests to perform poorly at predicting interactions (compared to predicting non-interactions). Reducing the number of non-interactions ('zeroes') by only including co-occurring pairs helps to balance the number of zeroes and non-zeroes in training data, and therefore likely improves the random forest models' predictive ability of interactions. In contrast to random forest, which uses interactions and non-interactions to make predictions, KNN only uses positive data (i.e. known interactions), thus avoiding the difficulty of making inferences from non-interactions (i.e., 'missing links' [38]), which could simply result from inadequate sampling.

#### KNN – adding 'new' host species

Because random forests use traits rather than species identities to predict interactions, interactions can be predicted between host and parasitoid species that did not occur in the training data. In contrast, KNN uses information on interactions among known hosts and parasitoids to predict interactions for new parasitoids (in the test data) with known hosts from the training data. To include host species that were only found in the test data, we used KNN to bring the new hosts into the training dataset by 'recommending' parasitoids from the training dataset to them, based on their nearest neighbours. For each new host species, we used host traits to find the k nearest host neighbours in the training data and took the average interaction frequency (in the training data) between the k host neighbours and each recommended parasitoid species (also from the training data) to calculate a predicted interaction frequency for each of the new host's interactions. We used the same method as described for parasitoids (see 'KNN' above) to calculate the Euclidean distance between host species, as all host traits were continuous, except for biogeographic status, which we encoded as a dummy variable and treated as a continuous trait. We made these predictions of parasitoids for new hosts rather than using the observed interaction frequency between the k host neighbours and their parasitoid species from the test data, because the latter approach would have essentially used the test data to train the model.

#### KNN – predicting interactions at test sites

To predict interactions at each test site using KNN, for each parasitoid species *i* at a site, we used species trait data to find the *k* nearest parasitoid species (in the training data) to parasitoid *i*.

We randomly shuffled the order of parasitoid species after each iteration to avoid the same species (those that arbitrarily appeared higher in the list) from being consistently preferred when their ranking was tied with others down the list. All host species that were attacked by these *k* parasitoid neighbours in the training data (and also present at the test site) were recommended to parasitoid *i* (i.e. the model predicts that parasitoid *i* will attack these host species). This is equivalent to recommending films that similar users have liked to a new user. To calculate a predicted interaction frequency for each of parasitoid *i*'s interactions, we took the average interaction frequency (in the training data) between the k neighbours and the recommended host species. This is different to random forests, which were trained on binary rather than weighted networks.

### Tuning hyperparameters

Random forest and KNN have a number of hyperparameters (including the number of trees in the forest and the number of neighbours, respectively) that can be tuned to optimise the model. Hyperparameters are configurations external to a model and their value cannot be learned by the model. Their value must be set by the user before the training process begins. In contrast, model parameters are learned during training (i.e. their value is estimated from the data) and are internal to the model.

The default hyperparameter settings for random forest and KNN are unlikely to be optimal for every given problem. Additionally, the default settings for random forest do not prevent a model from overfitting to the training data, and therefore may result in a model which performs well on the training data but poorly on the test data.

The best way to determine the optimal hyperparameter values is to try many different combinations and then assess the performance of each model. However, a complete grid search can be very computationally expensive if there are many hyperparameters (as in the case of random forest), since a model will be built for every possible combination of a pre-set list of values of the hyperparameters. For example, a grid with just 4 hyperparameters and 5 possible values for each hyperparameter gives $5^4 = 625$ possible combinations. By contrast, a random grid search, which does not try every combination but randomly selects combinations to try, provides a more efficient method and has been shown to almost always find the best hyperparameter settings [39]. We therefore used a random-grid search for random forest (as it has many hyperparameters) and a complete-grid search for KNN (which has fewer hyperparameters than random forest).

Hyperparameter tuning must be done on the training data. It is important that a model never 'sees' the test data until the testing phase (i.e. testing data should never be used in the training phase). However, optimising the hyperparameters for the training data can lead to overfitting (i.e. a model that performs well on the training data but poorly on the testing data). The standard way to account for this is to use cross validation [40]. K-fold cross validation (K-fold CV) is the most common cross validation method. K-Fold CV first splits the training set into K subsets (called 'folds'). The model is trained on K-1 of the folds and then evaluated on the Kth fold (called the validation data). This is repeated K times (i.e. for each fold). Performance metrics (calculated for each validation fold) are averaged to determine the final metrics for the model.

#### Tuning hyperparameters for random forest

To tune hyperparameters for the random forest, we used Scikit-learn's (v 0.23.1) 'RandomizedSearchCV' method, which uses a random grid search to select hyperparameter combinations (from a pre-defined list of values for each hyperparameter) and performs K-fold cross validation for each combination of values. Overall metric scores for all combinations are then compared to find the best hyperparameter combination.

Because our data are imbalanced (there are fewer interactions than non-interactions) and random forest uses both interaction occurrence and non-occurrence to train a model, we used stratified K-Fold CV; a variation of K-fold suitable for imbalanced data that returns stratified folds (i.e. the percentage of samples in each class is preserved). We used this instead of K-Fold, which randomly splits the data and does not necessarily preserve the percentage of samples in each class, because the small proportion of interactions ('ones') in the training data means that standard K-Fold may result in folds which only contain 'zeros' (i.e. non-interactions). A model trained on data that only contains 'non-interactions' would only be able to predict non-interactions.

Additionally, imbalance in the training data often causes the random forest classifier (and other classifiers) to perform poorly on the minority class (Jeni et al. 2013). For example, a binary matrix of parasitoids by hosts would have many more zeroes (where a given parasitoid does not attack a given host) than ones (where an interaction is present). To balance the training data and increase the classifier's ability to correctly predict the minority class (e.g. interaction occurrences), three main approaches are used: oversampling the minority class, under-sampling the majority class and a combination of both. Oversampling techniques can be random (for example, 'random oversampling of the minority class' in which random instances of the minority group are repeated) or synthetic (in which synthetic samples, similar to existing samples, are created). Similarly, under-sampling techniques can be random (e.g. random instances of the majority class are removed), but more complicated techniques exist in which, for example, observations that are near the border of the two classes are removed to help the classifier more easily distinguish between the two classes (e.g., 'Near-Miss' and Condensed Nearest Neighbour [41, 42]). Sampling methods are only performed on the training data.

Therefore, we performed a random grid search with stratified K-Fold CV to find the optimal hyperparameters for each random forest model ('native', 'plantation', 'combined') using Scikit-learn's

RandomizedSearchCV method with 100 iterations (i.e. hyperparameter combinations) and 3 folds (K=3). Further, we repeated this process for six different sampling methods (RandomOverSampler, RandomUnderSampler, NearMiss (version 1,2,3), and CondensedNearestNeighbour, all from the 'imblearn' v 0.6.2 package [43]) to find the best sampling method to balance the training data (see above paragraph). Only the training data were used to tune the hyperparameters and find the best sampling method.

#### Tuning hyperparameters for KNN

The number of neighbours, k, used to predict interaction partners is an important hyperparameter in KNN, and must be set before the model training process begins. It is often not possible to know the optimal number of neighbours to use prior to running the model. If parasitoid species interact with similar host species to their closest neighbours, a small value of k is likely to capture most interactions. However, if parasitoid species are more dissimilar in their interactions, a higher value of k is likely to better predict interactions, as a greater diversity of host species is likely to be recommended to a given parasitoid species. Additionally, neighbours can be unweighted (all neighbours contribute equally to predictions) or they can be weighted by distance (i.e. closer neighbours contribute more to predictions).

In contrast to random forests, where less-important predictors (traits) contribute less to predictions, in KNN all traits are equally weighted when determining the similarity between parasitoid species, and thus contribute equally to predictions. Furthermore, KNN assumes that similar parasitoid species (i.e. species with similar trait values) interact with a similar subset of host species. Therefore, it is important to select traits that make species with similar values for these traits likely to interact with a similar subset of host species (i.e. it is important to determine which traits correlate most strongly with similar interaction patterns), because the addition of uninformative traits will reduce predictive accuracy.

To determine the best parameter settings (k, whether to use weighted vs. unweighted neighbours, and which species traits to include) for each KNN model ('native', 'plantation', 'combined'), we performed a complete grid search (i.e. every subset of species traits with every value of k between 1 and 15, both weighted and unweighted) combined with leave-one-out cross validation on the training data. Specifically, for each parameter combination, we performed leave-one-out cross validation, in which each parasitoid species was iteratively removed from the training data, and the remaining training data were used to predict its interaction partners. For the runs with neighbour weightings, we used the weighting function described in Dudani [44], which weights neighbours inversely proportional to their distance from the focal parasitoid species (in trait-and-phylogenetic space). This use of a complete grid search combined with leave-one-out cross validation differs to random forest, where we performed a random grid search combined with stratified K-fold CV. We performed a complete grid search here for KNN because the small number of parameters made this computationally feasible and used leave-one-out cross validation to minimise cases of 'new' hosts (i.e., host species only found in the 'left-out' data, which occur when the parasitoid species left out is the only parasitoid species to interact with a given host species in the training data.).

#### Determining the best parameter combinations for random forest and KNN

To determine the best hyperparameter combination for each of the six models ('native', 'plantation' and 'combined' for both random forest and KNN), we calculated the F1 score (which only considers whether a prediction of an interaction occurrence is correct or incorrect) for each parameter combination. We chose this metric because it is appropriate for random forest (the F1 score is commonly used for imbalanced data [45] and our data are imbalanced), and because it can be used for KNN.

##### Random forest

The random forest classifier produces a probability of interaction occurrence for each possible parasitoid-host combination. Since the testing data include interactions that were either present or absent, and the F1 score requires a binary predicted presence/absence of interactions, it is necessary to set a threshold probability that is deemed to be an interaction presence. The default probability threshold for a random forest classifier is 0.5, though this may only be appropriate for balanced data (i.e. an equal number of presences and absences). However, this threshold can be adjusted to make the model more (or less) sensitive to false positives, and in the case of imbalanced data, in which the positive class (i.e. interactions; 'ones') is the minority, lowering the probability threshold is one method that can be used to increase a model's predictive ability of the positive class (though often at the cost of more false positives).

Ideally, a hyperparameter search would be crossed with a probability search (i.e. for every hyperparameter combination, the classifier's performance would be evaluated at all prescribed possible probability threshold values). However, the high computational cost of such a search made this unfeasible. We therefore first ran a hyperparameter search (described above) for each of the three models (i.e. 'native', 'plantation', 'combined') with a default probability threshold of 0.5, to find the best combination using the F1 score. Although the F1 score only considers whether a prediction of an interaction occurrence is correct or incorrect, and therefore depends on the threshold selected, the hyperparameter combination with the best F1 score also had the best average precision score (this was true for all three models). The average precision score (the area under the piecewise constant precision-recall curve) is a scoring rule – a method for evaluating the accuracy of a classifier based on predicted probabilities, rather than class labels, and is therefore independent of the threshold selected.

We then performed a search for the best probability threshold for each of the three random forest models, using the best hyperparameter settings found in the previous searches. Specifically, we performed stratified k-fold CV on the training data, in which for each iteration, we evaluated the classifier's performance at 19 equally spaced probability thresholds between 0.05 and 0.95, inclusive, and used the F1 score, as before, to determine the best probability threshold. Because the average precision score evaluates a classifier's performance based on predicted probabilities, and KNN produces predicted frequencies, rather than probabilities, we used the F1 score to determine the best parameter settings to be consistent across the two methods.

##### KNN

KNN predicts interaction frequencies, whereas the F1 score requires a binary predicted presence/absence of interactions (to assess its correctness). Therefore, to calculate the F1 score for each parameter combination, we converted each KNN predicted interaction frequency into a binary value. Because it is not possible to know *a priori* the best threshold for assigning the presence vs. absence of an interaction occurrence, we evaluated each parameter combination (for each model; 'native', 'plantation', 'combined') at 40 log (base 10) spaced thresholds between 0.05 and the largest predicted interaction frequency value. We used a log scale (rather than equally spaced points as for the random forest) because, unlike the random forest which predicts a probability, KNN predicts an interaction frequency, and using a log scale gives higher resolution to smaller numbers where the best threshold is more likely to be. We performed a search for the best threshold rather than assigning all non-zero predicted interaction frequencies as 1, as this would have resulted in a high number of false positives. In contrast to random forest, where we performed a probability search after the hyperparameter search, for KNN we performed the probability search for every possible hyperparameter combination to find the best settings. We used the same method described above (for recommending host species to parasitoid species) to select the best parameters for recommending parasitoid species to new host species.

#### Best hyperparameters for random forest

Random oversampling of the minority class (i.e. 'RandomOverSampler') was the best sampling method for all three random forest models, and the optimal probability threshold across the three models was 0.43. This is the average of the best probability thresholds for the three models (0.40, 0.40, and 0.50 for the 'native', 'plantation' and 'combined' models, respectively). The best hyperparameter values for the three random-forest models were: n_estimators=1366, min_samples_split=5, min_samples_leaf=5, max_features='log2', max_depth=13, criterion='entropy', class_weight='balanced'. Note that the best hyperparameter values were the same for all three random forest models ('native', 'plantation', 'combined'), and the names of the hyperparameters are as given in the Python 'RandomForestClassifier' function.

#### Best hyperparameters for KNN

For recommending host species to parasitoid species, and for recommending parasitoid species to new host species, weighting neighbours by distance (i.e. trait dissimilarity) gave a higher F1 score for 2/3 and 3/3 models, respectively, compared with using unweighted neighbours., We therefore used weighted neighbours for both hosts and parasitoids. The best value of k (number of neighbours) across the three models was 13 for recommending host species to parasitoid species and 11 for recommending parasitoid species to host species.

Across the weighted parasitoid neighbour models, all traits except phylogeny (i.e. body size, phenology, parasitoid ND, abundance) were collectively included in the 'native', 'plantation' and 'combined' models with the best F1 score. We therefore included these traits in the final models. This differs from random forests, which included all traits, but down-weighted less informative ones.

For recommending parasitoid species to host species (i.e. for including new host species only found in the test data), all host traits except abundance (i.e. body size, phylogeny, and phenology) were collectively included in the best 'native', 'plantation' and 'combined' models (with weighted neighbours). We therefore included these host traits in the final models.

The best overall threshold (average of the best threshold values for the three models: ‘native’, ‘plantation’, ‘combined’) for assigning presence vs. absence of an interaction occurrence was 0.19 for the host models (i.e. recommending parasitoids to hosts) and 0.14 for the parasitoid models.

The same parameter settings (including the probability threshold for converting a predicted probability/frequency into a binary value) were used for all three KNN models (‘native’, ‘plantation’, ‘combined’). Likewise, the same parameter settings were used across the three random-forest models. This decision was made because all traits were included in the random forest models (with less important traits down-weighted), whereas for KNN models important traits must first be selected. Therefore, we used the same parameter settings for all three KNN models to ensure the same traits were present in all the models, and we used the same parameter settings across the random-forest models to be consistent in our hyperparameter selection process across the two machine-learning techniques (and in our case the best hyperparameter settings, though not probability threshold, were the same for all three random forest models, see above).

### Model performance metrics

Class imbalance occurs naturally in a wide range of areas (e.g. ecological networks where there are few interactions compared to non-interactions), and while many solutions have been proposed, they mostly relate to either data resampling or model training, whereas the importance of selecting a suitable performance evaluation metric is often underestimated [46]. Accuracy (the proportion of correctly classified examples) is a commonly used metric for evaluating the performance of a classifier [47]. However, accuracy is only appropriate for data where the number of observations within each class is balanced, and can be misleading for imbalanced data [47]. For example, consider the extreme case of predicting a rare disease that occurs in 0.1% of the population. A model which classifies every individual as not having the disease will have an accuracy score of 99.9% but is clearly not useful.

In contrast to accuracy, the F1 score (the harmonic mean of precision and recall) is suitable for imbalanced data.

F1 score = (2 x precision x recall) / (precision + recall)

The F1 score varies between 0 (worst) and 1 (best). The harmonic mean (rather than average) is used because it penalises extreme values. For example, a classifier with a precision of 1 and a recall value of 0 will have an average of 0.5, but an F1 score of 0. Precision (also called positive predictive value) is the number of true positives divided by all positive predictions (i.e. precision = TP/(TP+FP)), and represents the proportion of correctly predicted interactions (‘ones’) from all host-parasitoid combinations that were predicted to interact. Low precision indicates a high number of false positives (e.g. host-parasitoid pairs that do not interact but are predicted to interact). Recall (also called sensitivity or the true positive rate) is the number of true positives divided by the number of positive values in the test data (i.e. recall = TP/(TP+FN)). Recall measures the proportion of correctly predicted interactions (‘one’) from all true interactions, and therefore captures a model’s ability to ‘recover’ all true interactions. High recall indicates a lower number of false negatives (e.g. host-parasitoid pairs that interact but are predicted to not interact). This is important in a biocontrol context, where the precautionary principle would suggest that it is better to predict false interactions than miss true ones. However, a model with only high recall, but low precision, would be of limited use in a biocontrol setting, as the high number of false positives would not allow useful prioritization of potential non-target host species for risk assessment. Therefore, an ideal model would have high recall and precision (and therefore a high F1 score), as precision and recall are complementary measures. For example, classifying all values as positive (negative) gives a recall (precision) of 1 but low precision (recall). Recall and precision both have a value of zero when ‘non-interacting’ (i.e. ‘zero’) is predicted for all host-parasitoid pairs, regardless of the number of true negatives (i.e. non-interactions; ‘zeros’).

Precision and recall are suitable metrics for imbalanced data because they do not use true negatives. In contrast, the false positive rate (i.e. FP / (FP + TN)) is not a suitable metric for imbalanced data because a significant increase in the number of false positives only increases the false positive rate by a small amount because the total number of negatives (FP+TN; the dominator) is large.

### Tests on the prediction of interaction occurrence

#### Method

To determine whether our ability to predict interaction occurrence (using machine-learning models) varies with species generality, we fitted a binomial generalised linear mixed effect model using predictive accuracy (i.e. whether a prediction of an interaction occurrence is correct or incorrect) as the response variable (Model set 1). As measures of generality, we included parasitoid and host normalised degree as interacting fixed effects. When making predictions about the impacts of novel species, such as potential biocontrol agents, it is possible that interaction and trait data will not exist for the specific recipient habitat. Therefore, we also included forest type (of the test site) as a non-interacting fixed effect to test whether predictions can be made equally within the same vs. across habitats. We included site and host-parasitoid pair as random factors (for all full models in the set) to control for the non-independence of interactions within a given site or involving particular species. To improve model convergence, we scaled and centred all continuous predictor variables, used the 'bobyqa' optimiser (from the lme4 R package [48]), and extended the maximum number of iterations.

We fitted a separate version of this model to the predictions from each random forest and KNN model ('native', 'plantation', 'combined' for both). We followed the same model selection process, and Bernoulli process to account for performing multiple statistical tests, as described in the Methods in the Main text.

Host and parasitoid ND were calculated from the training 'meta-networks' (see 'Training meta-networks' above). For the 'native' model, species ND calculated from the 'native' training meta-network was used to make predictions for both the plantation and native test sites, and analogously the 'plantation' training meta-network was used for the 'plantation' model. For the 'combined' model, habitat-type-specific ND values were used, i.e. ND values calculated using the 'native' training meta-network were used for the native test sites (and the 'plantation' training meta-network for plantation test sites). This approach allowed us to determine how well network and species data from one habitat type can predict interactions at a different habitat type.

The above models tested whether predictions for a given interaction were correct or incorrect, but did not distinguish false positives from false negatives. In a biocontrol context, distinguishing between false positives and negatives is important, as the precautionary principle would suggest that predicting false interactions (i.e. false positives) is better than missing true ones (i.e. false negatives). Therefore, as an additional test of whether the ability of a machine-learning model to predict networks differed when it was trained on data from the same vs. different habitat, we tested the performance of each random-forest and KNN model ('native', 'plantation', 'combined' in both cases) on each test site. For each test site, we calculated four metrics of model performance: F1 score, recall, precision, accuracy (see 'Model performance metrics' above). Importantly, these four metrics only consider whether a prediction of an interaction occurrence is correct or incorrect (rather than accounting for interaction frequencies).

We fitted a linear mixed model for each metric (the response) with model identity (combinations of KNN/random forest and native/plantation/combined) and forest type (of the test site) as interacting fixed effects. We included site as a random factor to control for the non-independence of interactions within a given site for all full models in this set. We used a binomial error for the recall, precision and accuracy models (since these metrics are proportions) and a gaussian error for the F1 score model.

For each linear mixed model, we selected the best-fitting model following the same model selection process as described in the Methods in the Main text. We used a Tukey's test (using the glht function in the multcomp v 1.4.13 package [49]) to test for pairwise differences in performance across models ('native', 'plantation', 'combined') and forest types. Overdispersion was not present in any of the binomial models, and the assumptions of normality and homoscedasticity were met for the gaussian models.

### Results

#### Predictive ability of interaction occurrences varied with generality of the interacting partners for random forest models, but not for KNN models

Predictive accuracy differed for specialists vs. generalists (measured as the number of interaction partners: normalised degree, ND) for random forest, but not KNN, models. Host and parasitoid ND significantly interacted in the 'combined' random-forest model, and although this interaction coefficient was positive, the negative host and parasitoid ND main effects meant that overall predictive accuracy declined (albeit sub-additively due to the interaction) with increasing host and parasitoid ND. Only host ND (and forest type; see below) were retained in the best-fitting 'plantation' and 'native' random-forest models, though host ND was only significant in the 'native' model (where it was negative), meaning that predictive ability declined with increasing host ND in this model (Table 11 in Appendix S1).

The fixed and random (site and host-parasitoid pair) effects both captured a high amount of variance in the 'native' and 'combined' random-forest models (native: $R_m^2$= 0.259, $R_c^2$=0.983, combined: $R_m^2$=0.365, $R_c^2$=0.879). For both models, most (<96%) of this random effect variance was captured by the host-parasitoid pair effect, indicating species idiosyncrasies in interaction preferences. In contrast, the fixed and random effects captured almost none of the variance in the 'plantation' model ($R_m^2$ and $R_c^2$ < 0.0001).

In contrast to random forest, predictive ability of interaction occurrences by KNN models did not vary with the generality of the interaction partners (parasitoid and host ND were not retained in any of the best-fitting KNN models, except for the 'combined' model where parasitoid ND was retained in the best-fitting model but was not significant). The host-parasitoid pair random effect was retained in all three best-fitting models though explained essentially none of the variance in predictive accuracy ($R_c^2$ <0.0001 for all three models).

#### *Predictive ability of interaction occurrences differed between forest types for random forest, but not KNN, models*

Predictions of interaction occurrences by 2/3 random-forest models ('native' and 'plantation') were significantly better at native sites than plantation sites (i.e. the forest-type fixed effect was retained and significant in both these models). In contrast, predictive accuracy did not differ between forest types (of the test sites) for any of the KNN models (the forest type fixed effect was not retained in any of these models) (Tables 11 and 12 in Appendix S1).

#### *Model performance metrics (F1 score, recall, precision, accuracy)*

The additional tests to separate out the influence of false positives and negatives (using four metrics of model performance: F1 score, recall, precision, accuracy) showed, overall, that predictions were independent of the habitat for which predictions were made and, to a lesser extent, the source of the training data (Table 14 and 15 in Appendix S1). For random forest two of the four metrics (recall and accuracy) depended on the source of the training data. The 'plantation' model gave higher recall scores than the 'native' model. However, the 'native' model gave higher accuracy scores than both the 'plantation' and 'combined' models. For KNN, the combined model gave higher recall scores than the plantation model, though the other three metrics (F1 score, precision, accuracy) did not depend on the source of the training data. There was a low probability of 2/4 and 1/4 metrics, respectively, differing across the random-forest and KNN models by chance (p=0.0016 and p=0.0274 calculated using the Bernoulli process).

Overall, random-forest and KNN models predicted interactions equally well at native and plantation test sites. The forest-type (of the test site) fixed effect was only retained in the best-fitting random-forest precision model and the best-fitting KNN accuracy model, though it was only significant in the second model (where it indicated that predictions at plantation sites were more accurate than predictions at native forest sites). The probability of 1/4 metrics differing significantly across forest-type for the KNN models by chance alone is quite high (p = 0.0274; calculated using the Bernoulli process), so these results could plausibly have arisen through type I error.

The fixed effects (model identity, and model identity and forest type, respectively) were strongly explanatory for the random-forest and KNN accuracy models (marginal $R^2$ = 0.44 and 0.197), but weakly explanatory for the remaining models. The site random effect was retained in the best-fitting KNN accuracy model and all best-fitting random forest models (except for precision) and captured a reasonable amount of variance (conditional $R^2$ >0.65 for all models).

### Discussion

#### Predictive ability of interaction occurrences varied with generality of the interacting partners for random forest models, but not for KNN models

Congruent to the finding for predictions of interaction frequency for both approaches, predictions of interaction occurrence by random-forest models varied with the generality (ND) of the interacting partners. In particular, predictive ability of interaction occurrences decreased with increasing ND of the interaction partners, consistent with previous research (as discussed in Discussion in the Main text).

In contrast to the findings for predictions of interaction frequency for both approaches, and predictions of interaction occurrence for random forest, predictions of interaction occurrence by KNN models did not depend on the generality (ND) of the interacting partners. This is surprising since there are several possible reasons for why predictions may be worse for generalist parasitoids. First, it is possible that similar generalists (i.e. generalists with similar trait values) overlap more in their host use than do specialists, simply because they interact with a greater number of host species. Because KNN uses the preferences of similar parasitoids to recommend hosts to a new parasitoid, a greater overlap in host use by similar (i.e. neighbouring) parasitoids, would likely lead to better predictions. Another reason for predictions being better for interactions among generalists than specialists is that specialist parasitoid species with similar trait values may be dissimilar in their host use (see Discussion in Main text), meaning that host recommendations from neighbouring specialists will likely not be useful.

#### Habitat type

Predictions of host-parasitoid interaction occurrences using machine-learning techniques were equally effective in both habitat types (plantation and native forest) for KNN, but not random-forest, models. However, when false positives were distinguished from false negatives (i.e., for the four-performance metrics that collectively captured a model's ability to predict true positives and negatives while avoiding false positives and negatives: accuracy, F1 score, precision and recall) (see 'Model performance metrics' above) no difference in predictive ability between the habitat types was present (this was true for both KNN and random forest). The exceptions to this were all three KNN models, which predicted binary interaction occurrence better at plantation than native-forest sites, though this was only true for one metric of model performance (accuracy), so may well have been a type I error.

Consistent with the findings for predictions of interaction frequencies (see Results in Main text), binary predictions of host-parasitoid interaction occurrences, overall, did not depend on the source (i.e., plantation, native forest, or both plantation and native forest) of the training data. This was true for 4/6 binomial GLM models and 2/4 and 3/4 metrics, respectively, for random forest and KNN.

This lack of an effect of data source is surprising, as plantation forests may capture a parasitoid's true host-preferences better than native forests (and we would therefore expect models trained on plantation data to be better) for the following reason. Plantation forests may be a more stressful environment than native forests, which may force parasitoids to rely on the subset of their maximal host range to which they are best adapted. In fact, previous research on the same system showed that the proportion of interactions with strong coevolutionary signal was higher in the plantation than in the native forest [28]. Similarly, a host-parasitoid network study showed that, as temperature increased, genetically similar parasitoids become more likely to attack genetically similar hosts (i.e., there was a stronger phylogenetic signal within species) [50]. Both these studies suggest that stress can induce a narrowing of tropic (phylogenetic) breadth (in the case of Lavandero and Tylianakis [50] congruence occurred between genotypes rather than species). Further, the lower structural complexity of plantation forests may increase the search efficiency of parasitoid species (e.g., [51, 52]). Both these characteristics may result in parasitoid species preferentially attacking their preferred hosts, and if the preferred interactions are the best phylogenetically or trait matched, this would make methods that incorporate phylogenetic or trait matching (e.g., random forest) trained on plantation rather than native forest data better at predicting interactions. In this way, network data from plantation forests may be expected to better capture parasitoids' true host preferences, than network data from more structurally complex habitats (e.g., native forests), which may consist of a core of 'stable' well-trait matched interactions, with a periphery of poorly-matched more opportunistic interactions. Although we may expect random-forest models trained on plantation rather than native forest data to perform better, we would not expect this for KNN as KNN does not incorporate trait or phylogenetic matching, and we did not find a consistent signal of this expectation in either machine-learning method.

### Indirect effects

#### Measure of shared parasitism

To calculate a quantitative measure of shared parasitoids, also known as the potential for apparent competition, $d_{ij}$, we used Equation 6 from Müller *et al.* [53]:

$$d_{ij} = \sum_{k=1}^{P} \left[ \frac{\alpha_{ik}}{\sum_{l=1}^{P} \alpha_{il}} \frac{\alpha_{jk}}{\sum_{m=1}^{H} \alpha_{mk}} \right] \qquad (1)$$

where α is the link strength (that is, number of attacks), i and j are a focal host species pair, m is all host species from 1 to H (the total number of host species), k is a parasitoid species, and l is all parasitoid species, from 1 to P (the total number of parasitoid species). When i=j, $d_{ij}$ represents the proportion of parasitoids attacking species i that recruit from species i. We included these cases to have a more complete 'picture' of the potential for apparent competition within the community, and because previous work on the same study system found that predictions of indirect effects did not differ qualitatively when such cases were removed [31].

### Calculating α from predicted networks

We calculated initial attack rates (α) for the random forest and KNN approaches from the predicted networks at time *t* test sites by the random-forest 'combined' model and the 'KNN' combined model, respectively. KNN produces a predicted interaction frequency for each host-parasitoid pair, so we calculated α directly from this. However, random forest produces a probability of interaction occurrence for each host-parasitoid pair. Therefore, to calculate α, for each host-parasitoid pair, using the random-forest networks, we divided the random-forest-predicted probability for each pair by the sum of predicted probabilities for all interactions involving the given parasitoid species (to calculate its proportional attack of each host species) and multiplied this value by the parasitoid abundance at the given site. We then summed these values for all the parasitoids of each host species to obtain its α value.

### Observed parasitism rate

$$O_{i(t+1)} = \frac{\sum_{l=1}^{P} \alpha_{il(t+1)}}{n_{i(t+1)}} \qquad (2)$$

where O is the observed parasitism rate of each host species *i* at time *t+1* test sites, and all other variables are defined as in equation 1 in 'Predicting indirect effects' in the Main text.

#### Tests using $d_{ij}$ calculated from predicted network data

##### Methods

To calculate the potential for apparent competition, $d_{ij}$, from the random forest and KNN predicted networks, we combined the predicted network for each test site into one ‘meta-web’, by taking the average random-forest-predicted probability (or KNN-predicted frequency) for each host-parasitoid pair. We included only cases where the pair co-occurred at a site, because non-co-occurring species may still be capable of interacting. We used the average value for each pair rather than summing values to avoid cases of probabilities greater than 1 for the random-forest-predicted ‘meta-web’. We then used these meta-webs to calculate the potential for apparent competition using the PAC function from the bipartite v 2.15 package [54]. These predicted ‘meta-webs’ contained non-integer values. However, the calculation for the potential for apparent competition is scale invariant (i.e. multiplying the entire interaction matrix by a common factor does not change the $d_{ij}$ values), meaning the relative potential for apparent competition can be calculated for interaction matrices containing non-integer values. We then used these $d_{ij}$ values to calculate expected parasitism rates (with α and host abundances being calculated in the same way as the analyses using observed rather than predicted networks to calculate $d_{ij}$), and also followed the same model fitting and selection processes.

##### Results

For these random-forest and KNN approaches, where predicted network data were used to calculate the potential for apparent competition ($d_{ij}$), expected parasitism rate was not retained in the best-fitting model for the random forest case, although it was significant (z value=2.35, p = 0.0186, df=78) in the KNN case and explained a reasonable amount of variance (marginal $R^2$ = 0.0963 (Table 18 and 19 in Appendix S1). However, the probability of this relationship being significant in 1/2 models by chance alone is moderately high (p=0.036; calculated using the Bernoulli process). The site random factor was retained in both models and captured a reasonable amount of variance (conditional $R^2$ =0.309, 0.343, respectively). Overall, for random forest approaches, these predictions were worse than with the other approach that used observed network data from the training dataset to calculate the potential for apparent competition, while for KNN approaches they were similar (Table 8 in Appendix S1).

### Coefficient tables for models in Main text

**Table 1. Predicting observed interaction frequency using random-forest models.** Results from the Poisson generalised linear mixed effect models showing the relationship between random-forest-predicted probability (scaled) (probability_sc) and observed interaction frequency. All models included parasitoid identity (paras_web_ID) as a random effect, and the 'native' and 'combined' models also included a random slope for predicted probability (probability_sc).

A) 'Native' model

```
Random effects:
 Groups       Name           Variance Std.Dev. Corr 
 paras_web_ID (Intercept)    0.9361   0.9675        
              probability_sc 0.5185   0.7201   -0.74
Number of obs: 671, groups:  paras_web_ID, 29

Fixed effects:
               Estimate Std. Error z value Pr(>|z|)    
(Intercept)     -2.4480     0.2527  -9.688  < 2e-16 ***
probability_sc   0.5792     0.2072   2.796  0.00518 ** 
---
```

B) 'Plantation' model

```
Random effects:
 Groups       Name        Variance Std.Dev.
 paras_web_ID (Intercept) 0.5359   0.732   
Number of obs: 671, groups:  paras_web_ID, 29

Fixed effects:
               Estimate Std. Error z value Pr(>|z|)    
(Intercept)     -2.5407     0.2207 -11.514  < 2e-16 ***
probability_sc   0.8629     0.1225   7.044 1.87e-12 ***
---
```

C) 'Combined' model

```
Random effects:
 Groups       Name           Variance Std.Dev. Corr 
 paras_web_ID (Intercept)    1.1527   1.0736        
              probability_sc 0.7849   0.8859   -0.90
Number of obs: 671, groups:  paras_web_ID, 29

Fixed effects:
               Estimate Std. Error z value Pr(>|z|)    
(Intercept)     -2.6476     0.2873  -9.216  < 2e-16 ***
probability_sc   0.8726     0.2373   3.677 0.000236 ***
---
```

**Table 2. Predicting observed interaction frequency using KNN models.** Results from the Poisson generalised linear mixed effect models showing the relationship between KNN-predicted frequency (scaled) (predicted_frequency_sc) and observed interaction frequency. All models included parasitoid identity (paras_web_ID) as a random effect, and the 'native' and 'combined' models also included a random slope for predicted frequency (predicted_frequency_sc).

A) 'Native' model

```
Random effects:
 Groups       Name                  Variance Std.Dev. Corr
 paras_web_ID (Intercept)           0.7710   0.8781
              predicted_frequency_sc 0.7514   0.8668   0.78
Number of obs: 671, groups:  paras_web_ID, 29

Fixed effects:
            Estimate Std. Error z value Pr(>|z|)
(Intercept)   -2.302      0.192  -11.99   <2e-16 ***
---
```

B) 'Plantation' model

```
Random effects:
 Groups       Name        Variance Std.Dev.
 paras_web_ID (Intercept) 0.5451   0.7383
Number of obs: 671, groups:  paras_web_ID, 29

Fixed effects:
                       Estimate Std. Error z value Pr(>|z|)
(Intercept)            -2.21839    0.19708  -11.26  < 2e-16 ***
predicted_frequency_sc  0.34574    0.07248    4.77 1.84e-06 ***
---
```

C) 'Combined' model

```
Random effects:
 Groups       Name                  Variance Std.Dev. Corr
 paras_web_ID (Intercept)           0.3097   0.5565
              predicted_frequency_sc 1.1630   1.0784   0.54
Number of obs: 671, groups:  paras_web_ID, 29

Fixed effects:
            Estimate Std. Error z value Pr(>|z|)
(Intercept)  -2.3924     0.1772   -13.5   <2e-16 ***
```

**Table 3.** AIC values and marginal and conditional $R^2$ values for Poisson generalised linear mixed effect models showing the relationship between random-forest-predicted probability (or KNN-predicted frequency) and observed interaction frequency for all three models ('native', 'plantation', 'combined') for both random-forest and KNN approaches.

| Model | AIC | $R_m^2$ | $R_c^2$ |
|---|---|---|---|
| Random forest 'native' | 528.07 | 0.058 | 0.312 |
| Random forest 'plantation' | 498.71 | 0.097 | 0.167 |
| Random forest 'combined' | 507.52 | 0.168 | 0.597 |
| KNN 'native' | 539.55 | 0 | 0.245 |
| KNN 'plantation' | 536.56 | 0.017 | 0.094 |
| KNN 'combined' | 500.85 | 0 | 0.219 |

**Table 4. Predictions of observed interaction frequency using random-forest models varied with the generality (measured as normalised degree; ND) of the interacting partners for 1/3 models.** Results from the Poisson generalised linear mixed effect models showing the effect of host and parasitoid generality (host_nd_sc and paras_nd_sc, respectively) on the relationship between random-forest predicted probability (probability_sc) and observed interaction frequency. All fixed effects were scaled and centred. Parasitoid identity (paras_web_ID) was included as a random effect in all three models, and a random slope for predicted probability (probability_sc) was included in the 'native' and 'combined' models.

A) 'Native' model

```
Random effects:
 Groups       Name           Variance Std.Dev. Corr
```

```
 paras_web_ID (Intercept)    0.7997   0.8943
              probability_sc 0.3985   0.6313   -0.79
Number of obs: 671, groups:  paras_web_ID, 29

Fixed effects:
                       Estimate Std. Error z value Pr(>|z|)
(Intercept)             -2.4921     0.2462 -10.121  < 2e-16 ***
probability_sc           0.7084     0.2381   2.975  0.00293 **
paras_nd_sc             -0.2901     0.1923  -1.509  0.13129
host_nd_sc              -0.1158     0.1495  -0.774  0.43864
paras_nd_sc:host_nd_sc   0.3535     0.1126   3.138  0.00170 **
---
```

B) 'Plantation' model

```
Random effects:
 Groups       Name        Variance Std.Dev.
 paras_web_ID (Intercept) 0.407    0.6379
Number of obs: 671, groups:  paras_web_ID, 29

Fixed effects:
                                      Estimate Std. Error z value Pr(>|z|)
(Intercept)                           -2.61239    0.24390 -10.711  < 2e-16 ***
probability_sc                         1.02014    0.17041   5.986 2.15e-09 ***
paras_nd_sc                           -0.56750    0.26388  -2.151  0.03151 *
host_nd_sc                            -0.37465    0.21460  -1.746  0.08085 .
probability_sc:paras_nd_sc             0.04271    0.18437   0.232  0.81679
probability_sc:host_nd_sc              0.02973    0.16143   0.184  0.85388
paras_nd_sc:host_nd_sc                -0.10420    0.23251  -0.448  0.65405
probability_sc:paras_nd_sc:host_nd_sc  0.41021    0.15719   2.610  0.00907 **
---
```

C) 'Combined' model

```
Random effects:
 Groups       Name           Variance Std.Dev. Corr
 paras_web_ID (Intercept)    0.9020   0.9498
              probability_sc 0.6721   0.8198   -0.85
Number of obs: 671, groups:  paras_web_ID, 29

Fixed effects:
                       Estimate Std. Error z value Pr(>|z|)
(Intercept)             -2.7239     0.2781  -9.793  < 2e-16 ***
probability_sc           1.3906     0.3270   4.253 2.11e-05 ***
paras_nd_sc             -0.4685     0.1981  -2.365   0.0181 *
host_nd_sc              -0.5021     0.2183  -2.300   0.0214 *
paras_nd_sc:host_nd_sc   0.2922     0.1480   1.974   0.0483 *
---
```

**Table 5. Predictions of observed interaction frequency using KNN models vary with the generality (measured as normalised degree; ND) of the interacting partners.** Results from the Poisson generalised linear mixed effect models showing the relationship between KNN-predicted frequency (predicted_frequency_sc) and observed interaction frequency, and how this is influenced by host and parasitoid normalised degree (host_nd_sc and paras_nd_sc, respectively). All fixed effects were scaled and centred, and all models included an observation-level random effect to deal with model overdispersion.

A) 'Native' model

```
Random effects:
 Groups Name        Variance Std.Dev.
 ob     (Intercept) 2.374    1.541
Number of obs: 671, groups:  ob, 671

Fixed effects:
                                             Estimate Std. Error z value Pr(>|z|)
(Intercept)                                   -2.5502     0.3526  -7.233 4.73e-13 ***
predicted_frequency_sc                         2.1818     0.6420   3.398 0.000678 ***
paras_nd_sc                                   -0.5614     0.2241  -2.505 0.012241 *
host_nd_sc                                     0.2599     0.2137   1.217 0.223791
predicted_frequency_sc:paras_nd_sc            -2.0501     0.7165  -2.861 0.004221 **
predicted_frequency_sc:host_nd_sc             -1.5500     0.3433  -4.515 6.34e-06 ***
paras_nd_sc:host_nd_sc                         0.6835     0.2625   2.604 0.009211 **
predicted_frequency_sc:paras_nd_sc:host_nd_sc  1.1691     0.3238   3.611 0.000305 ***
```

```
---
```

B) 'Plantation' model

```
Random effects:
 Groups Name        Variance Std.Dev.
 ob     (Intercept) 2.701    1.643
Number of obs: 671, groups:  ob, 671

Fixed effects:
                                              Estimate Std. Error z value Pr(>|z|)
(Intercept)                                   -3.26058    0.34837  -9.360  < 2e-16 ***
predicted_frequency_sc                         0.53817    0.18909   2.846 0.004425 **
paras_nd_sc                                   -0.04047    0.16443  -0.246 0.805604
host_nd_sc                                     0.39940    0.16815   2.375 0.017540 *
predicted_frequency_sc:paras_nd_sc            -0.46690    0.22519  -2.073 0.038143 *
predicted_frequency_sc:host_nd_sc             -0.66120    0.18571  -3.560 0.000370 ***
paras_nd_sc:host_nd_sc                         0.13794    0.17146   0.805 0.421100
predicted_frequency_sc:paras_nd_sc:host_nd_sc  0.48883    0.14542   3.362 0.000775 ***
---
```

C) 'Combined' model

```
Random effects:
 Groups Name        Variance Std.Dev.
 ob     (Intercept) 2.096    1.448
Number of obs: 671, groups:  ob, 671

Fixed effects:
                                              Estimate Std. Error z value Pr(>|z|)
(Intercept)                                    -2.8483     0.3415  -8.341  < 2e-16 ***
predicted_frequency_sc                          0.9924     0.2667   3.720 0.000199 ***
paras_nd_sc                                    -0.2039     0.1746  -1.168 0.242671
host_nd_sc                                      0.4673     0.1744   2.680 0.007354 **
predicted_frequency_sc:paras_nd_sc             -0.9379     0.3013  -3.113 0.001852 **
predicted_frequency_sc:host_nd_sc              -0.8879     0.1944  -4.567 4.94e-06 ***
paras_nd_sc:host_nd_sc                          0.4803     0.1920   2.502 0.012349 *
predicted_frequency_sc:paras_nd_sc:host_nd_sc   0.5788     0.1426   4.059 4.92e-05 ***
---
```

**Table 6.** Marginal and conditional $R^2$ values for the Poisson generalised linear mixed effect models showing the effect of host and parasitoid generality (measured as normalised degree; ND) on the relationship between random-forest-predicted probability (or KNN-predicted frequency) and observed interaction frequency for the three models ('native', 'plantation', 'combined') for both random-forest and KNN.

| Model | $R_m^2$ | $R_c^2$ |
|---|---|---|
| Random forest 'native' | 0.0981 | 0.3087 |
| Random forest 'plantation' | 0.1177 | 0.1709 |
| Random forest 'combined' | 0.2570 | 0.6019 |
| KNN 'native' | 0.1277 | 0.5922 |
| KNN 'plantation' | 0.0958 | 0.6062 |
| KNN 'combined' | 0.1317 | 0.5671 |

**Table 7. Predicting indirect effects using random forest and KNN approaches (with $d_{ij}$ values calculated from the observed training web)**. Results from the Binomial generalised linear mixed effect models showing the relationship between expected parasitism rate (all_E_sc) and observed parasitism rate, for both the random forest and KNN approach. Both models included site (block_trt_forest) as a random factor.

A) Random forest

```
Random effects:
 Groups           Name        Variance Std.Dev.
 block_trt_forest (Intercept) 0.3157   0.5619
Number of obs: 88, groups:  block_trt_forest, 29

Fixed effects:
            Estimate Std. Error z value Pr(>|z|)
```

```
(Intercept)  -2.0809     0.1582 -13.158   <2e-16 ***
all_E_sc      0.3840     0.1582   2.428   0.0152 *
---
```

B) KNN

```
Random effects:
 Groups           Name        Variance Std.Dev.
 block_trt_forest (Intercept) 0.2396   0.4895
Number of obs: 80, groups:  block_trt_forest, 29

Fixed effects:
            Estimate Std. Error z value Pr(>|z|)
(Intercept)  -2.1797     0.1510 -14.437  < 2e-16 ***
all_E_sc      0.3142     0.1107   2.839  0.00453 **
---
```

**Table 8.** Marginal and conditional $R^2$ values for models predicting indirect effects using random forest and KNN approaches (with $d_{ij}$ values calculated from the observed training web) (i.e. the Binomial generalised linear mixed effect models showing the relationship between expected parasitism rate and observed parasitism rate).

| Model | $R_m^2$ | $R_c^2$ |
|---|---|---|
| Random forest approach | 0.1222 | 0.3839 |
| KNN approach | 0.0962 | 0.3299 |

**Table 9.** AICc values and marginal and conditional $R^2$ values for the three approaches (random forest, KNN, data-based as used by Frost *et al.* [31]) for predicting indirect effects on a common data-set (N=20) (i.e. the models showing the relationship between expected parasitism rate and observed parasitism rate).

| Approach | AICc | $R_m^2$ | $R_c^2$ |
|---|---|---|---|
| Random forest | 63.10 | 0 | 0.344 |
| KNN | 63.10 | 0 | 0.344 |
| Data-based as used by Frost et al. 2016 | 63.10 | 0 | 0.344 |

**Table 10. Comparing the three approaches (random forest, KNN, data-based as used by Frost *et al.* [31]) for predicting indirect effects on a common data-set (N=20).** Results from the Binomial generalised linear mixed effect models showing the relationship between expected parasitism rate (using one of the three methods) and observed parasitism rate, for each of the three approaches. All models included site (block_trt_forest) as a random factor.

A) Random forest

```
Random effects:
 Groups           Name        Variance Std.Dev.
 block_trt_forest (Intercept) 0.2327   0.4824
Number of obs: 20, groups:  block_trt_forest, 17

Fixed effects:
            Estimate Std. Error z value Pr(>|z|)
(Intercept)  -1.6911     0.2286  -7.398 1.38e-13 ***
---
```

B) KNN

```
Random effects:
 Groups           Name        Variance Std.Dev.
 block_trt_forest (Intercept) 0.2327   0.4824
Number of obs: 20, groups:  block_trt_forest, 17

Fixed effects:
            Estimate Std. Error z value Pr(>|z|)
```

```
(Intercept)  -1.6911     0.2286  -7.398 1.38e-13 ***
---
```

C) Data-based approach as used by Frost *et al.* [31]

```
Random effects:
 Groups           Name        Variance Std.Dev.
 block_trt_forest (Intercept) 0.2327   0.4824
Number of obs: 20, groups:  site_trt_forest, 17

Fixed effects:
            Estimate Std. Error z value Pr(>|z|)
(Intercept)  -1.6911     0.2286  -7.398 1.38e-13 ***
---
```

### Coefficient tables for models in Appendix S1

**Table 11. Predicting interaction occurrence (i.e. binary interactions) using random forest models**. Results from the Binomial generalised linear mixed effect models showing the effect of host and parasitoid generality, as well as forest type (of the test site), on the ability of random-forest models ('native', 'plantation', and 'combined') to predict interaction occurrences. For all models, the response variable was predictive accuracy (i.e. whether a prediction of an interaction occurrence was correct or incorrect). All models included host-parasitoid pair as a random factor, and the 'native' and 'combined' models also included site as a random factor.

A) 'Native'

```
Random effects:
 Groups       Name        Variance Std.Dev.
 h_p          (Intercept) 174.308  13.203
 name_site_id (Intercept)   3.282   1.812
Number of obs: 671, groups:  h_p, 409; name_site_id, 29

Fixed effects:
            Estimate Std. Error z value Pr(>|z|)
(Intercept)   2.6817     1.0466   2.562   0.0104 *
host_nd_sc   -7.9290     1.2650  -6.268 3.66e-10 ***
forest_idP   -1.9053     0.9512  -2.003   0.0452 *
---
```

B) 'Plantation'

```
Random effects:
 Groups Name        Variance Std.Dev.
 h_p    (Intercept) 279.2    16.71
Number of obs: 671, groups:  h_p, 409

Fixed effects:
            Estimate Std. Error z value Pr(>|z|)
(Intercept)  -8.7244     0.7453 -11.706   <2e-16 ***
forest_idP   -1.1438     0.5228  -2.188   0.0287 *
host_nd_sc   -0.8540     0.5374  -1.589   0.1121
---
```

C) 'Combined'

```
Random effects:
 Groups       Name        Variance Std.Dev.
 h_p          (Intercept) 16.5807  4.0719
 name_site_id (Intercept)  0.6535  0.8084
Number of obs: 671, groups:  h_p, 409; name_site_id, 29

Fixed effects:
                      Estimate Std. Error z value Pr(>|z|)
(Intercept)            -1.5510     0.5000  -3.102  0.00192 **
host_nd_sc             -3.3125     0.8343  -3.970 7.18e-05 ***
paras_nd_sc            -0.4903     0.3595  -1.364  0.17257
host_nd_sc:paras_nd_sc   1.1330     0.4091   2.769  0.00562 **
---
```

**Table 12.** Predicting interaction occurrence (i.e. binary interactions) using KNN models. Results from the Binomial generalised linear mixed effect models showing the effect of host and parasitoid generality, as well as forest type (of the test site), on the ability of KNN models ('native', 'plantation', and 'combined') to predict interaction occurrences. For all models, the response variable was predictive accuracy (i.e. whether a prediction of an interaction occurrence was correct or incorrect). Host-parasitoid pair was included as a random factor in all models.

A) 'Native'

```
Random effects:
 Groups Name        Variance Std.Dev.
 h_p    (Intercept) 440.4    20.98
Number of obs: 671, groups:  h_p, 409

Fixed effects:
            Estimate Std. Error z value Pr(>|z|)
(Intercept)   -9.556      0.699  -13.67   <2e-16 ***
```

B) 'Plantation'

```
Random effects:
 Groups Name        Variance Std.Dev.
 h_p    (Intercept) 486      22.04
Number of obs: 671, groups:  h_p, 409

Fixed effects:
            Estimate Std. Error z value Pr(>|z|)
(Intercept)  -9.2755     0.6612  -14.03   <2e-16 ***
```

C) 'Combined'

```
Random effects:
 Groups Name        Variance Std.Dev.
 h_p    (Intercept) 395.3    19.88
Number of obs: 671, groups:  h_p, 409

Fixed effects:
            Estimate Std. Error z value Pr(>|z|)
(Intercept)  -9.1906     0.6973  -13.18   <2e-16 ***
paras_nd_sc  -0.7562     0.4529   -1.67    0.095 .
---
```

**Table 13.** Marginal and conditional $R^2$ values for random forest and KNN models for predicting interaction occurrence (i.e. binary interactions). In other words, the Binomial generalised linear mixed effect models showing the effect of host and parasitoid generality, as well as forest type (of the test site), on the ability of KNN and random forest models ('native', 'plantation', and 'combined', for both) to predict interaction occurrences. For all models, the response variable was predictive accuracy (i.e. whether a prediction of an interaction occurrence was correct or incorrect). Host-parasitoid pair was included as a random factor in all 6 models, and site was included as a random factor in the 'native' and 'combined' random-forest models.

| Model | $R_m^2$ | $R_c^2$ |
|---|---|---|
| Random forest 'native' | 0.2592 | 0.9836 |
| Random forest 'plantation' | <0.0001 | <0.0001 |
| Random forest 'combined' | 0.3654 | 0.8797 |
| KNN 'native' | 0 | <0.0001 |
| KNN 'plantation' | 0 | <0.0001 |
| KNN 'combined' | <0.0001 | <0.0001 |

**Table 14.** Results for the linear mixed models assessing the performance of random forest for predicting binary interactions (using four metrics of model performance: F1 score, Accuracy, Recall, Precision). These metrics were the response variables, and we included model identity (i.e. 'native', 'plantation', 'combined') and forest type (of the test site) as predictor variables. We used a binomial error for the recall, precision and accuracy models and a gaussian error for the F1 score model. Site was included as a random factor in all models, except the model in which precision was the response variable.

A) F1 score

```
Random effects:
 Groups   Name        Variance Std.Dev.
 sites    (Intercept) 0.016883 0.12994
 Residual             0.005061 0.07114
Number of obs: 87, groups:  sites, 29

Fixed effects:
            Estimate Std. Error t value
(Intercept)  0.20288    0.02531   8.017
```

B) Accuracy

```
Simultaneous Tests for General Linear Hypotheses

Multiple Comparisons of Means: Tukey Contrasts

Fit: glmer(formula = y_accur ~ model_id + (1 | sites), data = data_CF,
    family = binomial)

Linear Hypotheses:
                         Estimate Std. Error z value Pr(>|z|)
Native - Both == 0         0.4711     0.1134   4.155   <1e-04 ***
Plantation - Both == 0    -0.8168     0.1242  -6.577   <1e-04 ***
Plantation - Native == 0  -1.2878     0.1234 -10.436   <1e-04 ***
---
```

C) Recall

```
Simultaneous Tests for General Linear Hypotheses

Multiple Comparisons of Means: Tukey Contrasts

Fit: glmer(formula = y_recall ~ model_id + (1 | sites), data = data_CF,
    family = binomial)

Linear Hypotheses:
                         Estimate Std. Error z value Pr(>|z|)
Native - Both == 0        -0.6120     0.5617  -1.090   0.5166
Plantation - Both == 0     1.4511     0.7596   1.910   0.1334
Plantation - Native == 0   2.0631     0.7578   2.723   0.0173 *
---
```

D) Precision

```
Simultaneous Tests for General Linear Hypotheses

Multiple Comparisons of Means: Tukey Contrasts

Fit: glm(formula = y_precision ~ forest_id, family = binomial, data = data_CF)

Linear Hypotheses:
           Estimate Std. Error z value Pr(>|z|)
P - N == 0  -0.2959     0.1651  -1.792   0.0731 .
---
```

**Table 15.** Results for the linear mixed models assessing the performance of KNN for predicting binary interactions (using four metrics of model performance: F1 score, Accuracy, Recall, Precision). These metrics were the response variables, and we included model identity (i.e. 'native', 'plantation', 'combined') and forest type (of the test site) as predictor variables. We used a binomial error for the recall, precision and accuracy models and a gaussian error for the F1 score model. Site was only included as a random factor in the accuracy model.

A) F1 score

```
Simultaneous Tests for General Linear Hypotheses

Multiple Comparisons of Means: Tukey Contrasts

Fit: lm(formula = f1_score ~ model_id, data = data_CF)

Linear Hypotheses:
                         Estimate Std. Error t value Pr(>|t|)
Native - Both == 0        0.01324    0.06480   0.204    0.977
Plantation - Both == 0   -0.11580    0.06480  -1.787    0.180
Plantation - Native == 0 -0.12905    0.06480  -1.991    0.121
```

B) Accuracy

```
Simultaneous Tests for General Linear Hypotheses

Multiple Comparisons of Means: Tukey Contrasts

Fit: glmer(formula = y_accur ~ forest_id + model_id + (1 | site_id),
    data = data_CF, family = binomial)

Linear Hypotheses:
           Estimate Std. Error z value Pr(>|z|)
P - N == 0   0.7765     0.2879   2.697    0.007 **
---
```

C) Recall

```
Simultaneous Tests for General Linear Hypotheses

Multiple Comparisons of Means: Tukey Contrasts

Fit: glm(formula = y_recall ~ model_id, family = binomial, data = data_CF)

Linear Hypotheses:
                         Estimate Std. Error z value Pr(>|z|)
Native - Both == 0        -0.2479     0.3526  -0.703   0.7615
Plantation - Both == 0    -1.0184     0.3648  -2.792   0.0148 *
Plantation - Native == 0  -0.7704     0.3632  -2.121   0.0855 .
---
```

D) Precision

```
Coefficients:
            Estimate Std. Error z value Pr(>|z|)
(Intercept)  -1.3063     0.1182  -11.05   <2e-16 ***
---
```

**Table 16.** Marginal and conditional $R^2$ values for linear mixed models assessing the performance of random forest for predicting binary interactions (using four metrics of model performance: F1 score, Accuracy, Recall, Precision). We used a binomial error for the recall, precision and accuracy models and a gaussian error for the F1 score model. Site was included as a random factor in all models, except the precision model. The best-fitting precision model was a GLM (the value listed under $R_m^2$ is the $R^2$ value).

| Model | $R_m^2$ | $R_c^2$ |
|---|---|---|
| F1 score | 0 | 0.7693 |
| Accuracy | 0.4434 | 0.7130 |
| Recall | 0.0864 | 0.6546 |
| Precision | 0.0774 | - |

**Table 17.** Marginal and conditional $R^2$ values for linear mixed models assessing the performance of KNN for predicting binary interactions (using four metrics of model performance: F1 score, Accuracy, Recall, Precision). We used a binomial error for the recall, precision and accuracy models and a gaussian error for the F1 score model. Site was only included as a random factor in the accuracy model.

| Model | $R_m^2$ | $R_c^2$ |
|---|---|---|
| F1 score | 0.0528 | - |
| Accuracy | 0.1970 | 0.7200 |
| Recall | 0.0959 | - |
| Precision | 0 | - |

**Table 18. Predicting indirect effects using random forest and KNN approaches (with $d_{ij}$ values calculated from the machine-learning predicted webs)**. Results from the Binomial generalised linear mixed effect models showing the relationship between expected parasitism rate (all_E_sc) and observed parasitism rate, for both the random forest and KNN approach. Both models included site (block_trt_forest) as a random factor.

A) Random forest

```
Random effects:
 Groups           Name        Variance Std.Dev.
 block_trt_forest (Intercept) 0.3216   0.5671
Number of obs: 85, groups:  block_trt_forest, 29

Fixed effects:
            Estimate Std. Error z value Pr(>|z|)
(Intercept)   -2.137      0.157  -13.61   <2e-16 ***
---
```

B) KNN

```
Random effects:
 Groups           Name        Variance Std.Dev.
 block_trt_forest (Intercept) 0.257    0.507
Number of obs: 81, groups:  block_trt_forest, 29

Fixed effects:
            Estimate Std. Error z value Pr(>|z|)
(Intercept)  -2.0672     0.1492 -13.857   <2e-16 ***
all_E_sc      0.3167     0.1345   2.354   0.0186 *
---
```

**Table 19.** Marginal and conditional $R^2$ values for models predicting indirect effects using random forest and KNN approaches (with $d_{ij}$ values calculated from the predicted web) (i.e. the Binomial generalised linear mixed effect models showing the relationship between expected parasitism rate and observed parasitism rate).

| Approach | $R_m^2$ | $R_c^2$ |
|---|---|---|
| Random forest | 0 | 0.3096 |
| KNN | 0.0963 | 0.3432 |

**Table 20.** Importance of (a) parasitoid- and (b) host- traits and phylogenies in random-forest models ('combined', 'native', and 'plantation'). To calculate the importance of each trait, and the importance of phylogenies, we used the 'feature_importances_' function from Python's v 0.23.1 Scikit-learn package [35], which uses Gini importance (also known as mean decrease impurity) to calculate the importance of each feature (i.e. trait or phylogeny).

A) Parasitoid Traits and Phylogeny

| Parasitoid trait or phylogeny | 'Combined' model | 'Native' model | 'Plantation' model | Average importance |
|---|---|---|---|---|
| Abundance | 0.213242 | 0.228813 | 0.239655 | 0.227237 |
| Phenology | 0.095252 | 0.087126 | 0.08286 | 0.088413 |
| Generality (ND) | 0.072078 | 0.072504 | 0.065541 | 0.070041 |
| Phylogeny | 0.052664 | 0.053643 | 0.058443 | 0.054917 |
| Body size | 0.022885 | 0.018939 | 0.016397 | 0.019407 |

B) Host Traits and Phylogeny

| Host trait or phylogeny | 'Combined' model | 'Native' model | 'Plantation' model | Average importance |
|---|---|---|---|---|
| Phenology | 0.09922 | 0.089587 | 0.114662 | 0.101156 |
| Phylogeny | 0.103787 | 0.090301 | 0.094277 | 0.096122 |
| Body size | 0.077761 | 0.078629 | 0.089155 | 0.081849 |
| Abundance | 0.080352 | 0.0848 | 0.067389 | 0.077514 |
| Generality (ND) | 0.058552 | 0.066407 | 0.060494 | 0.061818 |
| Biogeographic status | 0.052358 | 0.058413 | 0.043579 | 0.05145 |
